# Effects of n-Hexane and ethanol extract of *Milicia excelsa* stone on pain perception in Sodium Monoiodoacetate-induced Osteoarthritic Female Wistar Rats

**DOI:** 10.1101/2024.07.03.601862

**Authors:** Samson Adegbe, Lasaki Esohe Abiodun, Hidaayah Oluwamayowa Jimoh-Abdulghaffaar, Meekness Adegoke

**Affiliations:** Redeemer’s University; Altinbas University; University of Ilorin

**Keywords:** Osteoarthritis, *Milisia excelsa* stone, Pain biomarkers

## Abstract

**AIM:** This research was undertaken to evaluate the effect of n-Hexane and ethanol extract of *Milicia excelsa* stone on pain perception in Sodium Monoiodoacetate-induced Osteoarthritic Female Wistar Rats. And the most active ingredient in *Milicia excelsa* stone is calcium carbonate.

**METHOD:** For this study, fifty (50) female Wistar rats divided into five groups were used, these include: Positive control (normal health rats), Negative control (osteoarthritic rats with no treatment), Reference (osteoathritic rats treated with arthocare), Test (1) (osteoarthritic rats treated with n-Hexane extract of *milicia excelsa* stone), Test (2) (osteoarthritic rats treated with ethanol extract of *milicia excelsa* stone). Von Frey Hair Filament was used to access pain perception across the groups. And the knee circumference was taken pre-induction, induction, and post-induction.

**RESULTS:** The result of this experimental research showed that there was significant (p<0.05) decrease both in the cartilage and serum level of biomarkers such as Prostanglandin E_2_, Tumour Necrotic factor alpha, Vascular endothelial growth factor, and Cartillage oligomeric metalloprotein. There was significance (p<0.05) decrease in the knee oedema. Also, the pain threshold increased in both n-Hexane and ethanol groups.

**CONCLUSION:** It is therefore concluded that n-Hexane and ethanol extracts of *Milicia excelsa* have significant effect in reducing pain perception in monosodium iodoacetate-induced osteoarthritic female Wistar rats.

## INTRODUCTION

Osteoarthritis (OA) is a complex disease involving the whole synovial joint. It is the most common type of arthritis and a leading cause of disability due to pain. The most susceptible joint is the knee joint and it is common in older adults particularly women.

It was originally believed to occur due to wear and tear of the articular surface in joints. But current understanding points to a more complex process. Osteoarthritis involves degeneration of cartilage, abnormal bone remodeling, osteophyte formation and joint inflammation. The four major parts of the synovial joint that participate in this pathology are the meniscus (majority of synovial joints), articular cartilage, subchondral bone, and synovial membrane. These components in healthy joints provide support for the joint. The meniscus provides several functions including load bearing and shock absorption in the knee joint. It is a fibrocartilage composed mainly of water, type I collagen, and proteoglycans (predominantly aggrecan) in its extracellular matrix. Other components include type II, III, V, and VI collagen. The articular cartilage provides a surface for movement of the synovial joint. It is a hyaline cartilage composed mainly of proteoglycans and type II collagen in the matrix. It is divided into deep, middle, and superficial zones characterized by the differences in the matrix composition and cell orientation. Calcified cartilage serves as an interface between the bone and articular cartilage. The subchondral bone gives support to the joint and is composed of mineralized type I collagen. The synovial membrane (synovium) produces the synovial fluid. This fluid, which is composed of lubricin and hyaluronic acid, lubricates the joint and nourishes the articular cartilage. The synovium is composed of two types of synoviocytes: fibroblasts and macrophages. The synovial fibroblasts produce the synovial fluid components. The synovial macrophages are usually dormant but are activated during inflammation (Kuyinu *et al.,* 2016).

Several abnormalities to these components contribute largely to osteoarthritis. Mechanical abrasion in the knee can lead to the progressive degenerative changes in the meniscus with loss of both type I and, more severely, type II collagen. This effect initially occurs from the middle substance of each meniscus rather than the articulating surface. More importantly, recent studies point to an inflammatory mechanism for the initial stages of the disease. This occurs mainly in response to injury caused by mechanical stimulation of the joint. The release of cytokines, such as interleukin-1 (IL-1), IL-4, IL-9, IL-13, and TNF-α, degradative enzymes such as a disintegrin and metalloproteinase thrombospondin-like motifs (ADAMTS), and collagenases/matrix metalloproteinases (MMPs) by chondrocytes, osteoblasts, and synoviocytes triggers the process. Furthermore, the innate immune system plays a role in Osteoarthritis progression through the activation of both the complement and alternative pathways (Nair *et al.,* 2016).

Other factors may contribute to Osteoarthritis pathogenesis in the cartilage. In aging individuals, chondrocytes increase their production of inflammatory cytokines. Advanced glycation end products have also been implicated in this process. These AGEs accumulate in the articular cartilage in older individuals. They bind to receptors on chondrocytes leading to the release of proinflammatory cytokines and VEGF, ultimately leading to cartilage degeneration on this pathway illustrates the influence of age in the development of Osteoarthritis and endorses a sequence of natural disease occurrence. Adipokines, cytokines secreted by adipose tissue and the infrapatellar fat pad in the knee, have been linked with the degradation of articular cartilage. This implies the potential role of obesity, in the development of Osteoarthritis. Importantly, systemic inflammation has been posited as an additional pathologic feature of Osteoarthritis. Although many studies question if it plays a role in the disease process, due to the belief that Osteoarthritis is a focal disease, quite a few published works in recent years indicate that Osteoarthritis should be classified as a systemic musculoskeletal disease. The current findings on Osteoarthritis pathogenesis present cytokines and inflammation as possible targets of treatment. These could warrant the use of drugs against proinflammatory cytokines, such as anti-rheumatic drugs, in the treatment of the disease. These drugs have shown varying success in preclinical studies; however, they have not been fully tested in clinical studies. In addition to these, lifestyle modifications and other treatment methods may play important roles in the treatment and prevention of the disease (Narayana *et al.,* 2016).

### Classification of osteoarthritis

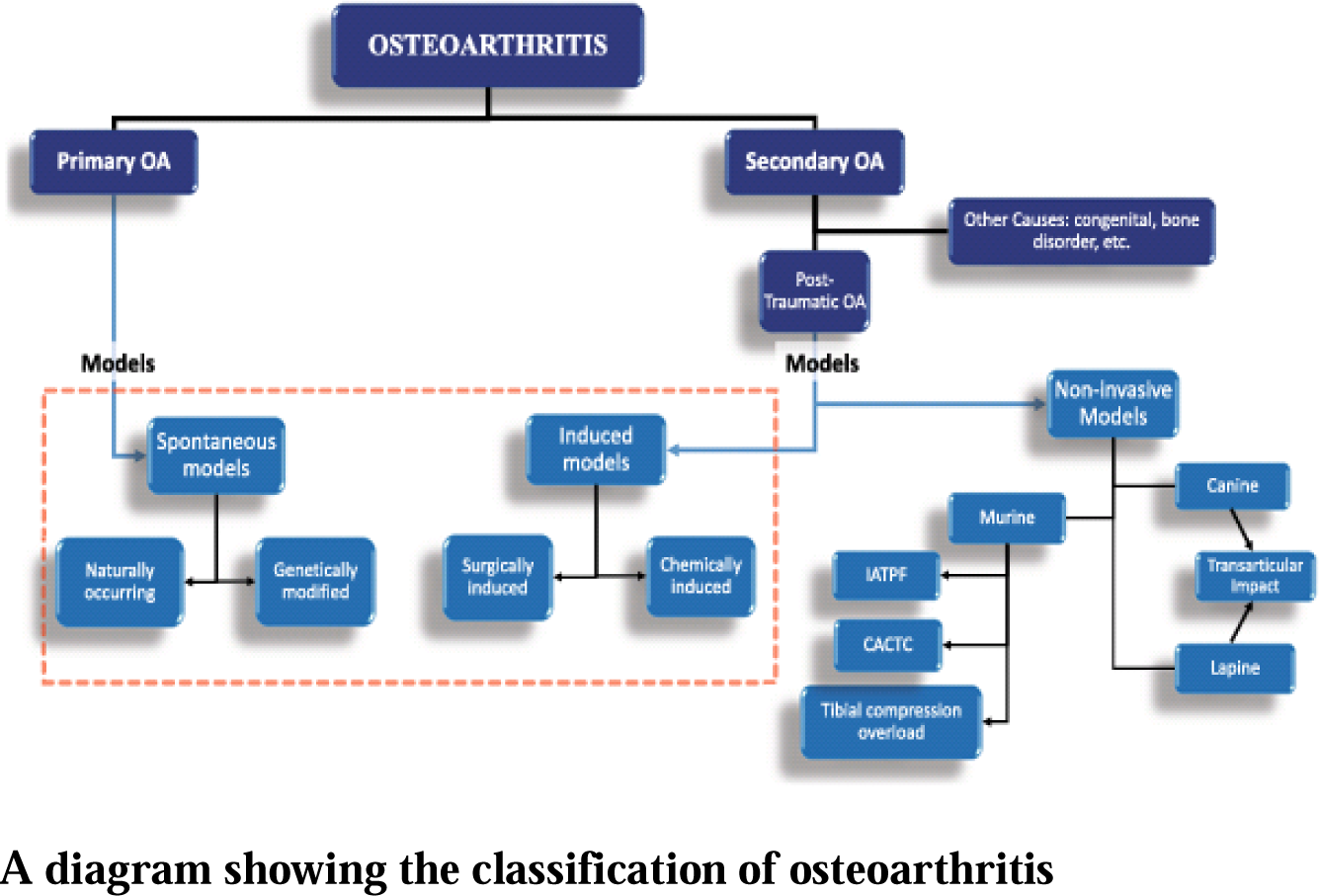

Osteoarthritis has typically been classified into primary (idiopathic) and secondary Osteoarthritis based on the disease etiology. Primary osteoarthritis (POA) is a naturally occurring phenomenon due to degenerative changes in the joint. It is further classified into localized and generalized Osteoarthritis. Localized OA affects one joint while generalized Osteoarthritis affects three or more joints. Secondary Osteoarthritis is normally associated with causes and/or risk factors leading to Osteoarthritis in the joint. These include trauma, congenital diseases, and other diseases or disorders of metabolism or the bone. It is important to note that the heterogeneous nature of Osteoarthritis presents challenges to its classification and treatment. For that reason, one treatment cannot apply to all patients with the disease. The variability of etiology, treatment, and outcomes for each patient makes the need to classify Osteoarthritis into clinical phenotypes a highly discussed venture. These discussions propose that categorizing Osteoarthritis into clinical phenotypes, adapted to their specific treatment, will improve patient outcomes. Based on these recommendations, five phenotypes have been proposed which replace the original primary and secondary classifications with features of the disease. These include post-traumatic, metabolic, aging, genetic, and pain phenotypes (Bendele *et al.,* 2001).

The post-traumatic Osteoarthritis phenotype is analogous to post-traumatic osteoarthritis (PTOA), which is caused by acute or repetitive injury to the joint. Patients with this phenotype would benefit from preventative measures, such as the use of braces in athletes, prevention from falls in older adults, and prevention of surgical intervention such as meniscectomies. The metabolic/obesity phenotype represents both the effect of increased loading on weight-bearing joints from obesity and the role of adipokines on the development of OA. Understanding this phenotype would help in therapy decisions such as exercise programs for weight loss goals and hormone therapy for menopause-related OA. The aging phenotype is most analogous to POA. It is a naturally occurring phenotype due to advanced aging of the individual. This phenotype could benefit from targeted therapy designed to inhibit AGEs and the cytokines released from senescent chondrocytes. The genetic phenotype is related to how hereditary factors affect the development of OA through complex mechanisms. These findings could provide specific targets for gene or drug therapy. Finally, the pain phenotype describes the development of OA pain due to inflammation and abnormal bone remodeling in the joint. The development of anti-inflammatory and pain medications would benefit patients in this phenotype. Although other clinical phenotypes have been described this proposal serves as the closest classification to understand the pathogenesis of the disease and its correlation to the animal models. These five phenotypes may also prompt increased discussion of the disease as we make new discoveries on its pathophysiology (Malfait *et al.,* 2015).

### MILISIA EXCELSA STONE

The frequent occurrence of large stone-like deposits in Iroko tree has been recognized for a long time now; doubt still appears to exist concerning the true composition and origin of the material. The Iroko stone is gotten from a log of Iroko tree cut or broken (Campbell *et al.,* 1932).

It is made of both organic and inorganic components. The inorganic component is made up of calcium carbonate, while the organic component is made up of wood borers, ants, which have calcified. It also includes wood chips, and bark (Record, 1927).

For the purpose of this study, the inorganic component of the stone was extracted using n-Hexane and ethanol. Below is a specimen of the Iroko stone.

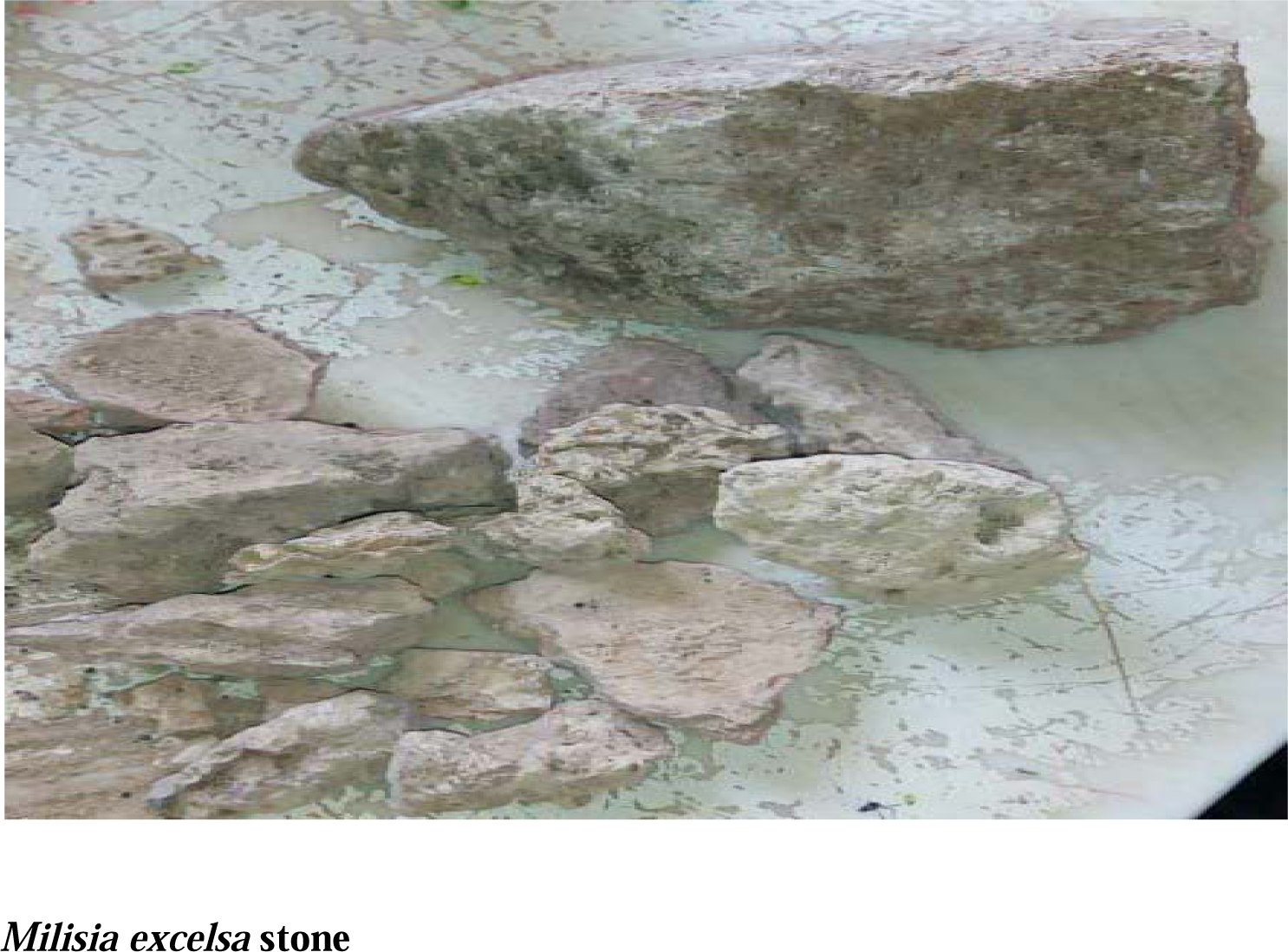

The model used for this study was chemical model. Monosodium idiodoacetate (MIA) was injected intra-articularly into the right knees of the female Wistar rats. And the biomarkers tested for were Prostaglandin E2 (PGE2), Tissue Necrotic Factor alpha (TNF-α) Vascular endothelial growth factor (VEGF), and Cartillage oligomeric metalloprotein (COMP). Mechanical withdrawal threshold of the hind paw was tested using Von Frey monofilament hair (VFH). VFHs with bending forces of 2-100g was applied to the center of the plantar region of the ipsilateral hind paw for 5 seconds, beginning with the lowest force, and hind paw withdrawal threshold was determined.

## METHODS

### METHODOLOGY

#### Experimental Design

This study was a comparative study among normal healthy rats, arthocare-treated rats, untreated osteoarthritic rats, *Milicia excelsa* Ethanol extract-treated osteoarthritic rats as well as *Milicia excelsa* n-Hexane extract-treated rats. It was carried out using 50 female albino rats (wistar rats) within the weight range of 180 – 250g. The animals were kept in the animal house situated in the University of Ilorin, College of Health Sciences (COHS). They were housed under natural, normal light and dark cycles at room temperature with the appropriate ventilation and spacing provided by the College. The plastic cages in which they were kept, together with the saw dust were carefully cleaned and replaced daily.

They were acclimatised for a period of two (2) weeks and were fed with standard rat pellet and distilled water *ad rem*. After acclimatization, they were injected with monoiodoacetate salt (MIA), to induce ostheoarthristis within a period of 12 days. Thereafter, the administration of *Milicia excelsa* for three weeks.

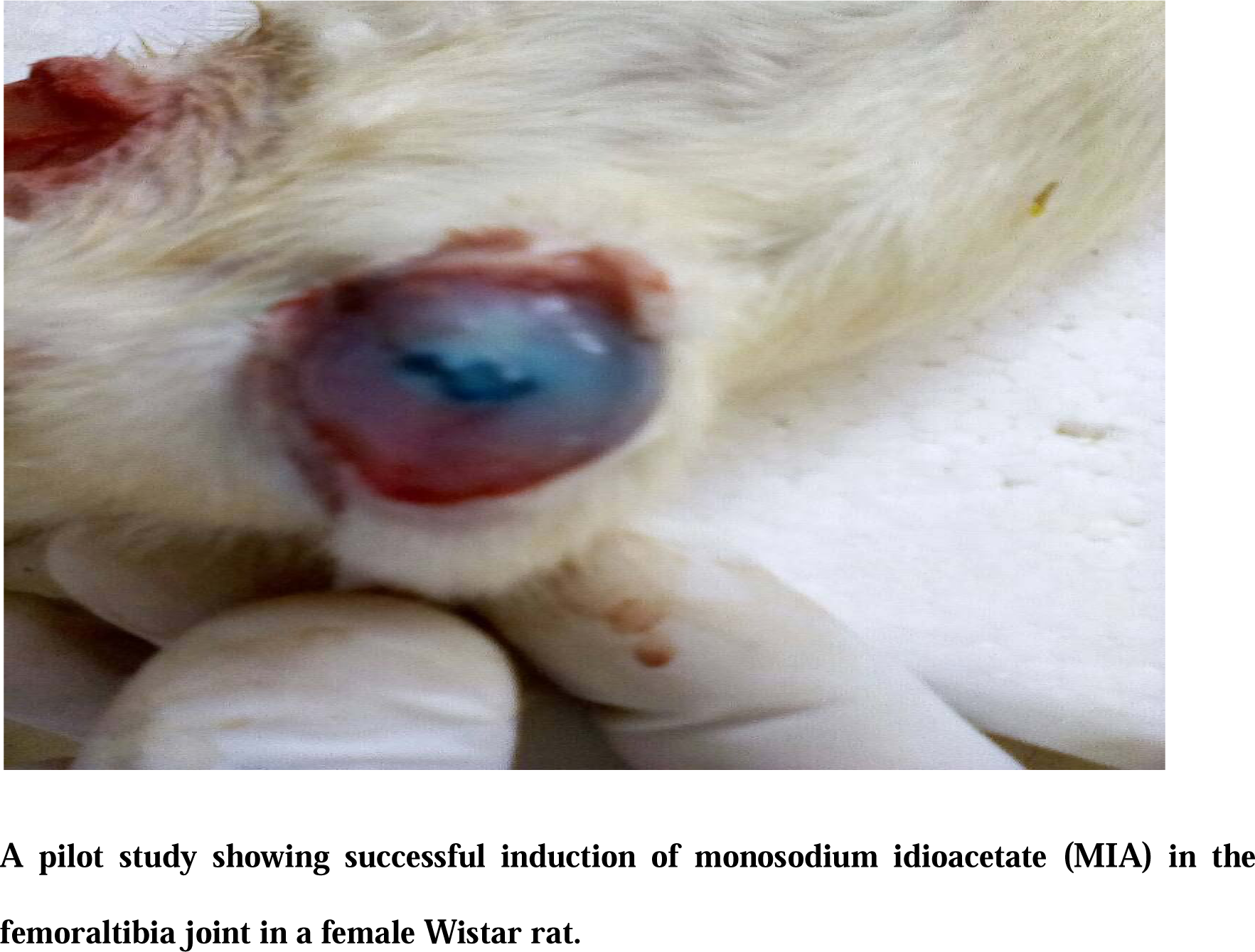

#### Processing of plant materials

*Milicia excelsa* stone (Iroko stone) was the only plant material used in this research. It was obtained from Lagos, Nigeria. The stone was pounded into powdery form using ceramic mortar and pestle, air dried for three (3) days before soaking in both ethanol and n-Hexane.

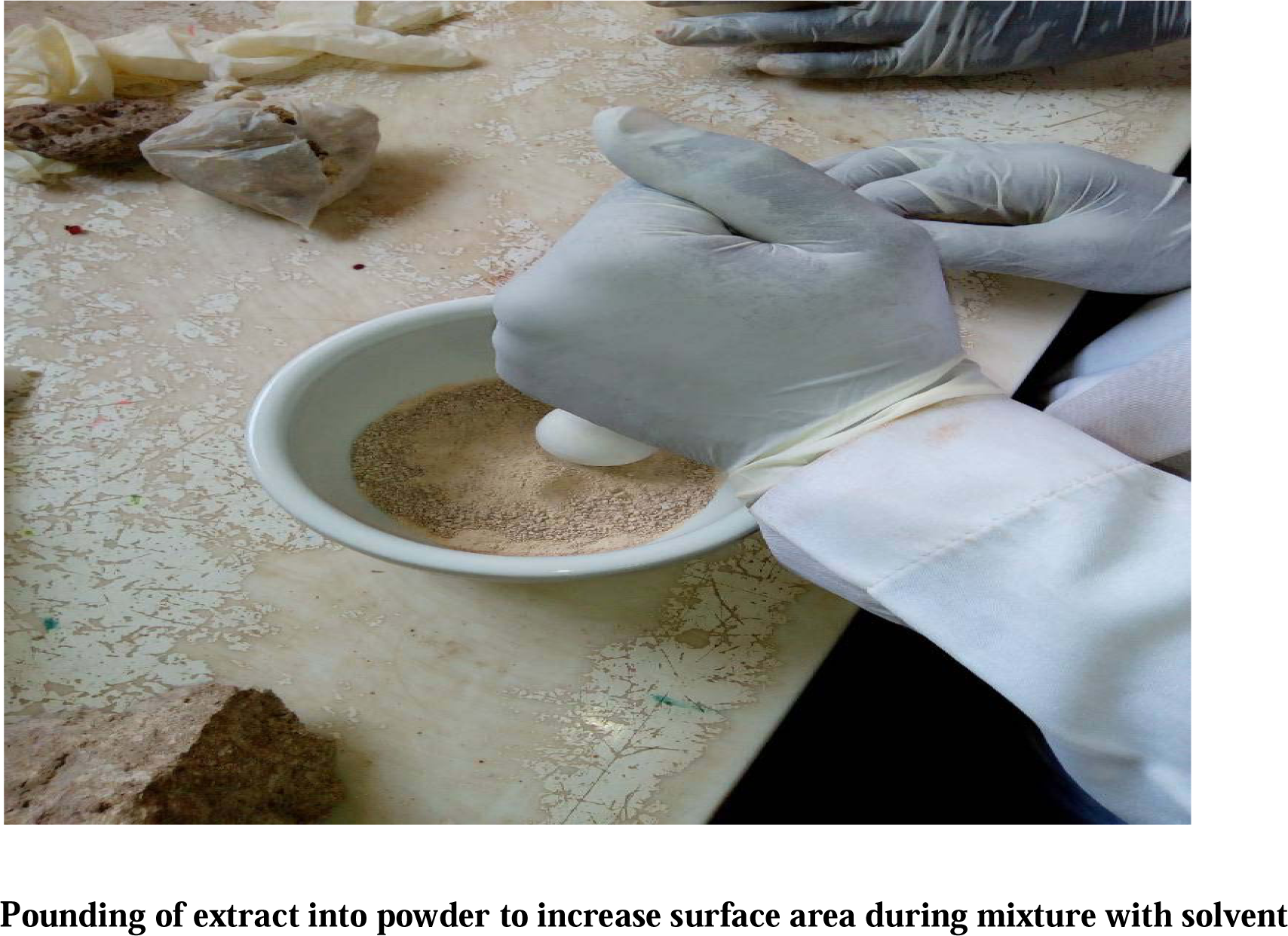

The processed Iroko stone was then divided into two, which were soaked in ethanol and n-Hexane respectively for three days. The mixtures were constantly stirred so that most of the content of the Iroko stone dissolved into the media. The fluid parts of these mixtures were then extracted by filtration using filter paper as the filter medium. The filtrate was afterwards poured into clean beaker and taken for centrifugation at the Central Research Laboratory, University of Ilorin.

The extract was evaporated using a Rotatary evaporator, evaporated filtrate was the dissolved in normal saline for daily administration.

### GROUPING AND ADMINISTRATION

The animals were randomly assigned into five (5) groups each containing ten (10) animals. The following tables show the distribution of the animals as in; their labels, size and treatment:

Distribution and grouping of Wistar Rats
| Group 1:<br>Control group |  |  |  |
| --- | --- | --- | --- |
| S/N | LABEL | SIZE | TREATMENT |
| 1-10 | 1 | 10 | 1mg of normal saline |

| Group 2: Osteoarthritis group |  |  |  |
| --- | --- | --- | --- |
| S/N | LABEL | SIZE | TREATMENT |
| 11-20 | 2 | 10 | 1mg of normal saline |

| Group 3: Arthocare group |  |  |  |
| --- | --- | --- | --- |
| S/N | LABEL | SIZE | TREATMENT |
| 21-30 | 3 | 10 | Osteoarthritis + Arthocare (0.00833g of chondritin sulphate and Glucosamine 0.00667g) |

| Group 4: n-Hexane group |  |  |  |
| --- | --- | --- | --- |
| S/N | LABEL | SIZE | TREATMENT |
| 31-40 | 4 | 10 | Osteoarthritis + 1000mg/kg of *Milisia excelsa* (Ethanoic extract) |

| Group 5: Ethanol group |  |  |  |
| --- | --- | --- | --- |
| S/N | LABEL | SIZE | TREATMENT |
| 41-50 | 5 | 10 | Osteoarthritis + 1000mg/kg of *Milisia excelsa* (n-Hexane extract) |

#### Method of administration

The rats were administered the ethanoic and n-Hexane *Milisia excelsa* extract through the oral route administration with the aid of an oral cannula according to their weight in grams from the stock solution already prepared.

#### Assessment of tactile allodynia (VON FREY)

Mechanical withdrawal threshold of the hind paw was tested using Von Frey monofilament hair (VFH). VFHs with bending forces of 2-100g was applied to the center of the plantar region of the ipsilateral hind paw for 5 seconds, beginning with the lowest force, and hind paw withdrawal threshold was determined. If the animal did not respond to the force of the VFH, the next higher force was applied until recoil is observed. Ascending method of force application was carried out four times post preliminary threshold and used to calculate the actual paw withdrawal time (PWT).

#### Knee edema measurement

Assessment of knee odema of animals was carried out with a thread to note if there were swellings due to pain as a result of the destruction of the chondrocytes by the monoiododacetate salt. This was also done before induction of the salt in order to have a baseline. It was then performed two hours after induction of the salts. Subsequent measurement thereafter was done on weekly basis.

#### Radiographic evaluation

The right induced knees of the animals were exposed to X-rays before and after treatment. Radiographs were made in craniocaudal position at full extension after anaesthesia with usual development process. The images were analysed by a radiologist without knowledge of joint and subgroup of animals evaluated.

#### Histopathology

After radiographic evaluation, the animals were sacrificed under a mild dose of ketamin hydrochloride and their knees were excised for histopathology.

### SAMPLE COLLECTION

At the end of the study period, the animals were sacrificed under a mild dose of ketamin. Blood was collected through cardiac puncture and their right knees were excised for histopathology. The blood collected was further centrifuged and the plasma was decanted for analysis. While the knees excised were weighed, and then homogenized in Phosphate Buffer Saline (PBS). Homogenates were then centrifuged at 3000rpm for 15 minutes and the resulting supernatants were stored in ice for biochemical analysis.

## RESULTS

The levels of Tumor Necrotic Factor, Prostanglandin E2, Vascular endothelial growth factor (VEGF), and Cartillage oligomeric metalloprotein (COMP) in n-Hexane and ethanol groups were compared with other groups. Also, their knee oedema and mechanical sensitivity to pain.

### BIOMARKERS

#### SERUM LEVEL ANALYSIS

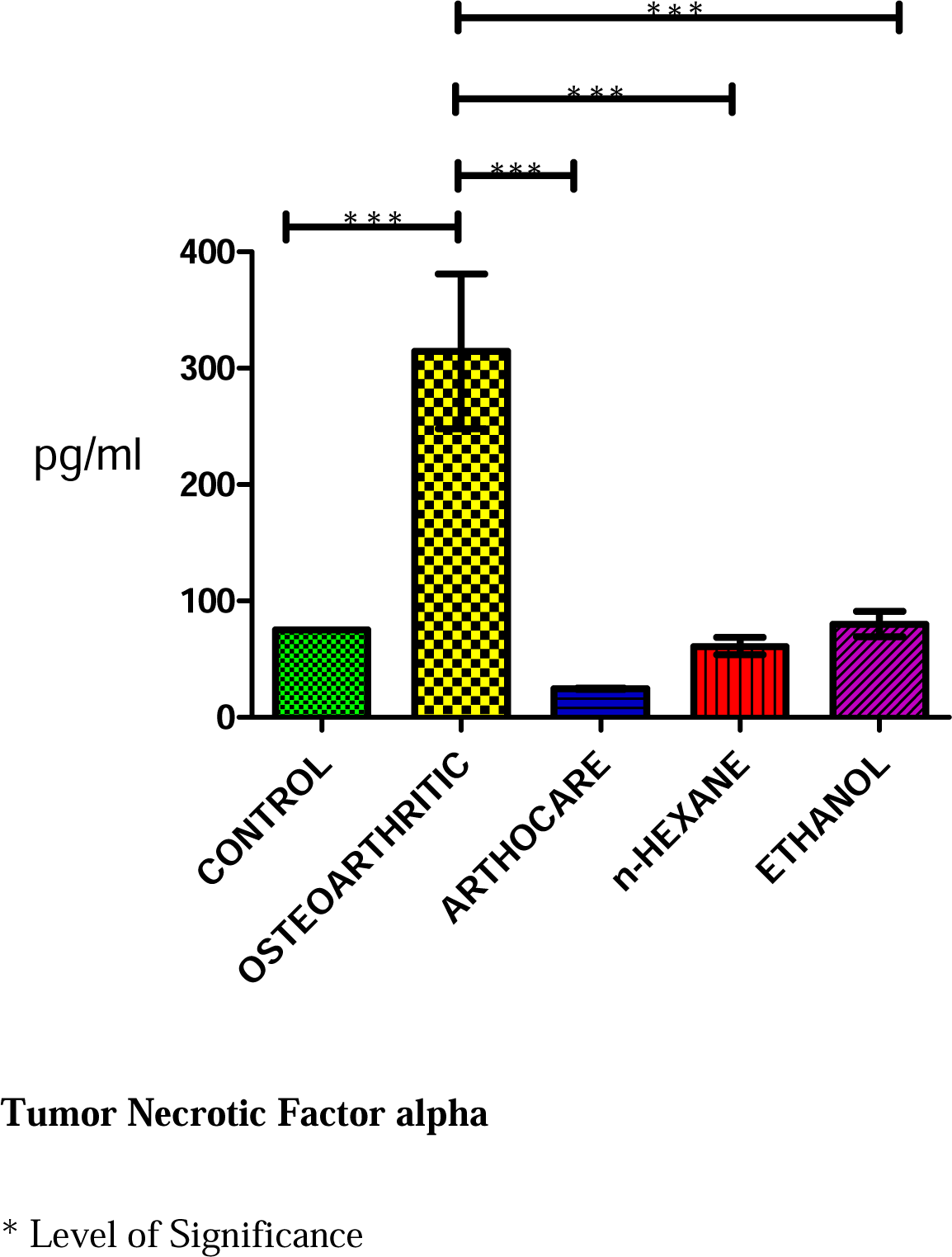

The level of TNF-α in n-Hexane group showed no significant difference when compared with the control group. It indicated marked significant (p<0.05) decrease when compared with the Osteoarthritic group. There was no significant (p<0.05) difference when n-Hexane group was compared with Arthocare and Ethanol group.

Also, the level of TNF-α in Ethanol group showed no significant (p<0.05) difference when compared with the control group. It indicated marked significant (p<0.05) decrease when compared with the Osteoarthritic group. There was no significant (p<0.05) difference when n-Hexane group was compared with Arthocare and Ethanol group.

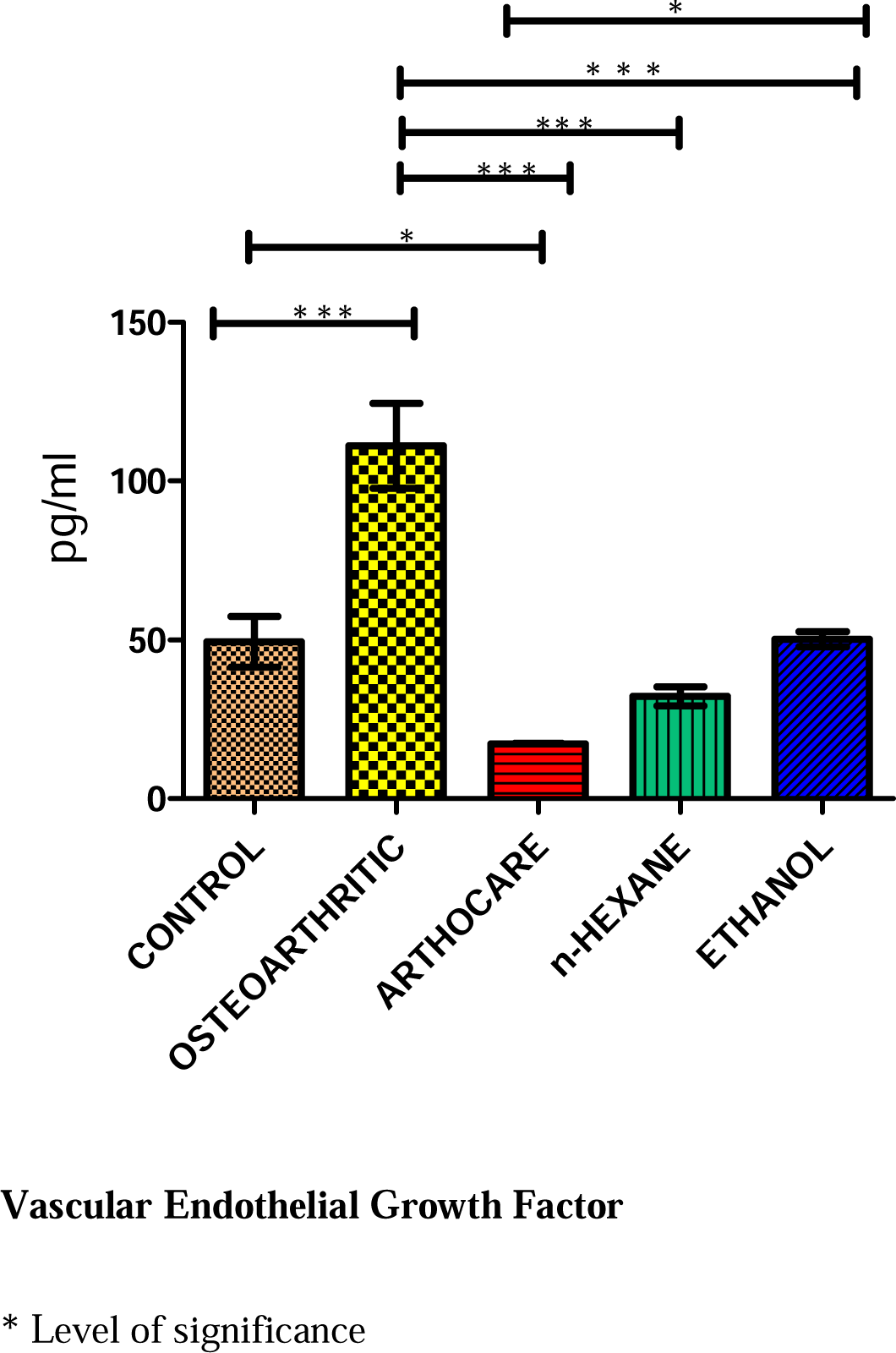

The level of VEGF in the serum showed no significant (p<0.05) decrease when n-Hexane group was compared with control group. There was significant (p<0.05) decrease when n-Hexane group was compared with osteoarthritic group. There was no significant (p<0.05) decrease when n-Hexane group was compared with both arthocare and ethanol groups respectively.

The level of VEGF in the serum showed no significant (p<0.05) decrease when ethanol group was compared with control group. There was significant (p<0.05) decrease when ethanol group was compared with osteoarthritic group. There was no significant (p<0.05) decrease when ethanol group was compared with both arthocare and n-Hexane groups respectively.

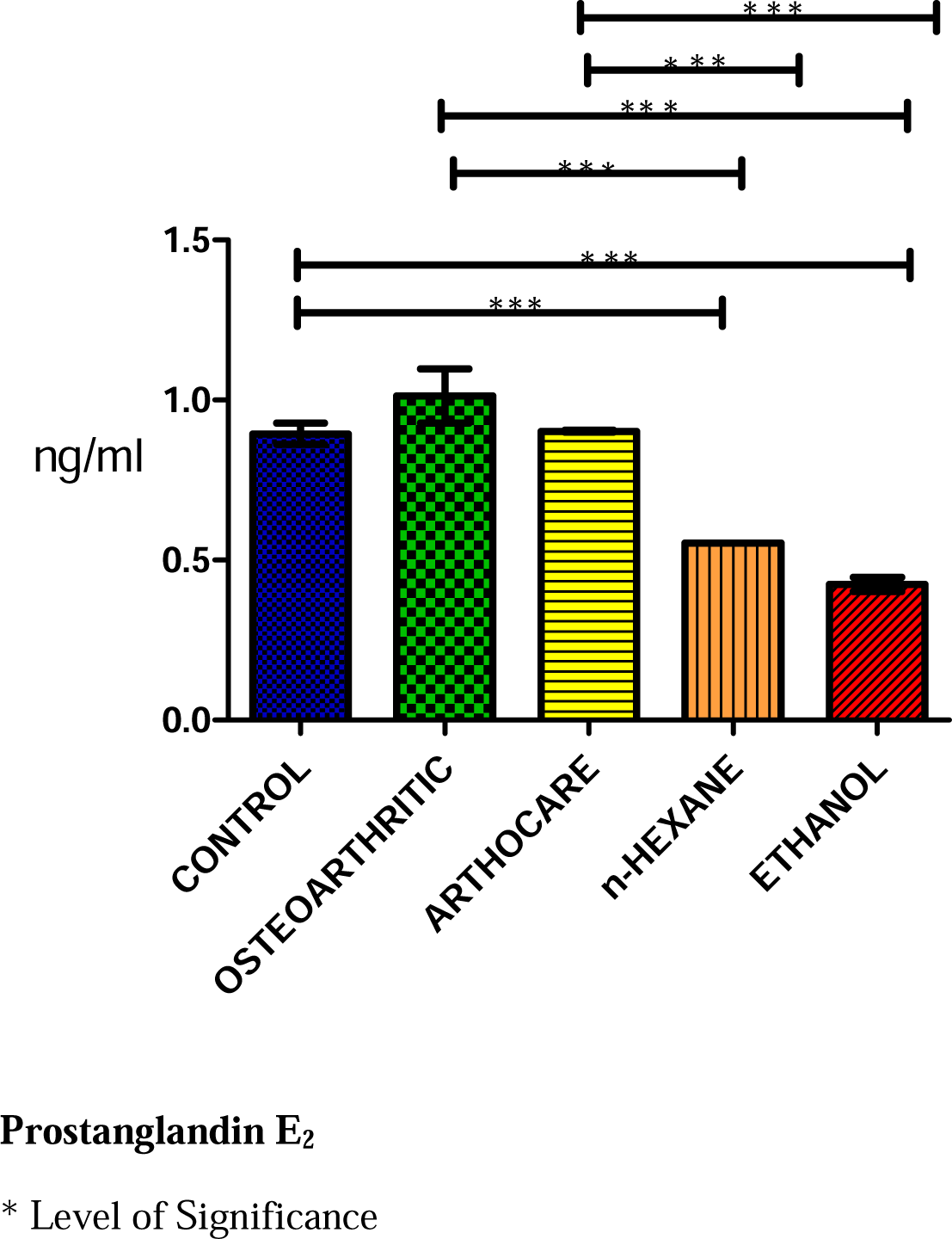

The level of Prostanglandin E_2_ in n-Hexane group showed significant (p<0.05) decrease when compared with the control group. It indicated marked significant (p<0.05) decrease when compared with the Osteoarthritic group. There was significant (p<0.05) decrease when n-Hexane group was compared with Arthocare group but no significant (p<0.05) difference when n-Hexane group was compared with Ethanol group.

Also, the level of Prostanglandin E_2_ in Ethanol group showed significant (p<0.05) decrease when compared with the control group. It indicated marked significant (p<0.05) decrease when compared with the Osteoarthritic group. There was significant (p<0.05) decrease when n-Hexane group was compared with Arthocare but no significant (p<0.05) difference when n-Hexane group was compared Ethanol group.

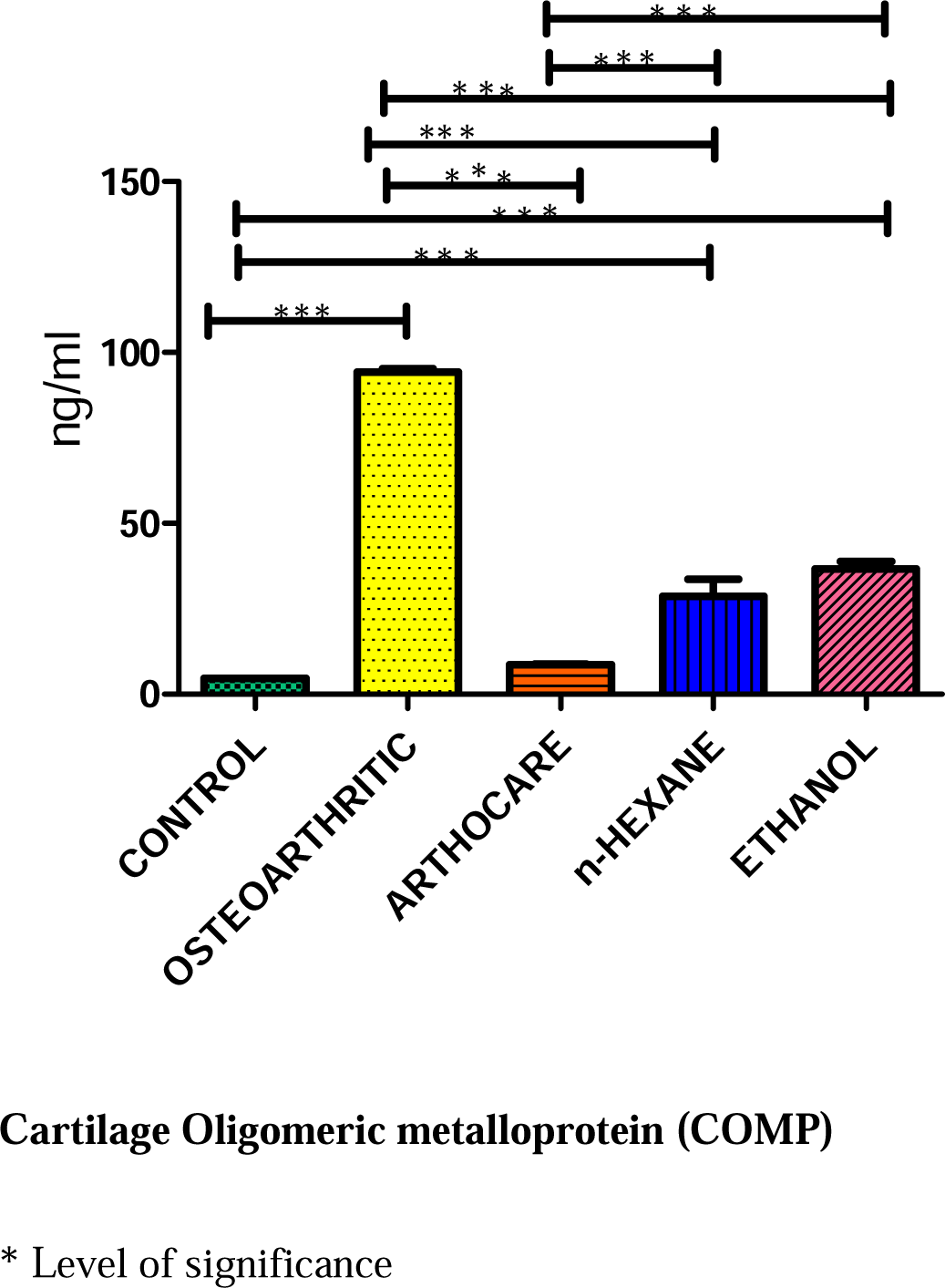

The level of COMP in the plasma showed significant (p<0.05) increase when n-Hexane group was compared with control group. There was significant (p<0.05) decrease when n-Hexane group was compared with osteoarthritic group. The level of COMP in the plasma showed significant (p<0.05) increase when n-Hexane group was compared with arthocare group. There was no significant (p<0.05) decrease when n-Hexane group was compared with ethanol group.

The level of COMP in the plasma showed significant (p<0.05) increase when ethanol group was compared with control group. There was significant (p<0.05) decrease when ethanol group was compared with osteoarthritic group. The level of COMP in the plasma showed significant (p<0.05) increase when ethanol group was compared with arthocare group. There was no significant (p<0.05) decrease when ethanol group was compared with n-Hexane group.

#### CARTILAGE LEVEL ANALYSIS

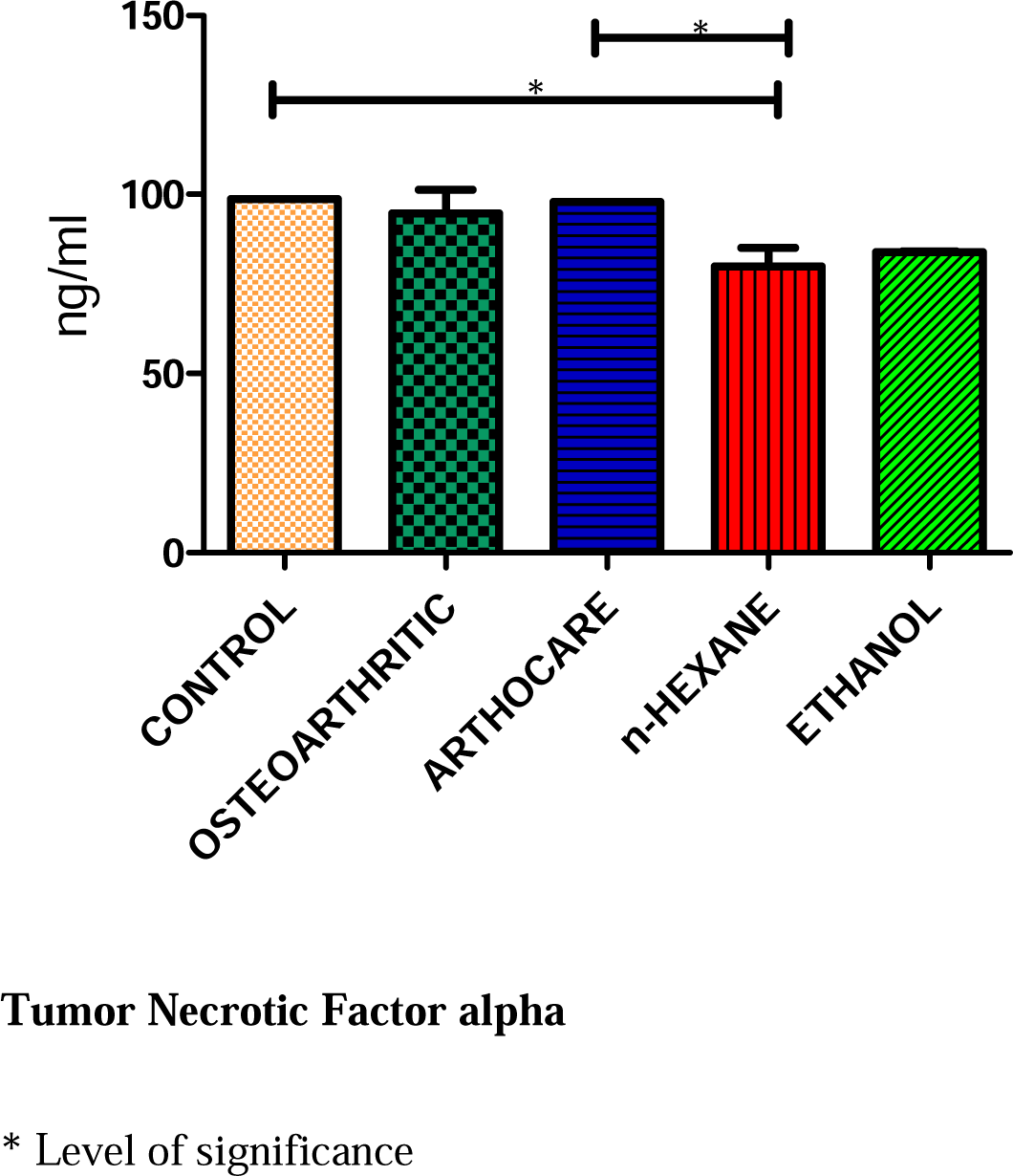

The level of TNF-α cartilage in tissue showed significant (p<0.05) decrease when n-Hexane group was compared with control group. There was no significant (p<0.05) decrease when n-Hexane group was compared with Osteoarthritic group. There was significant (p<0.05) decrease when n-Hexane group was compared with Arthocare group. There was no significant (p<0.05) decrease when n-Hexane group was compared with Ethanol group.

The level of TNF-α cartilage in tissue showed no significant (p<0.05) decrease when Ethanol group was compared with control group, osteoarthritic, arthocare and n-Hexane groups.

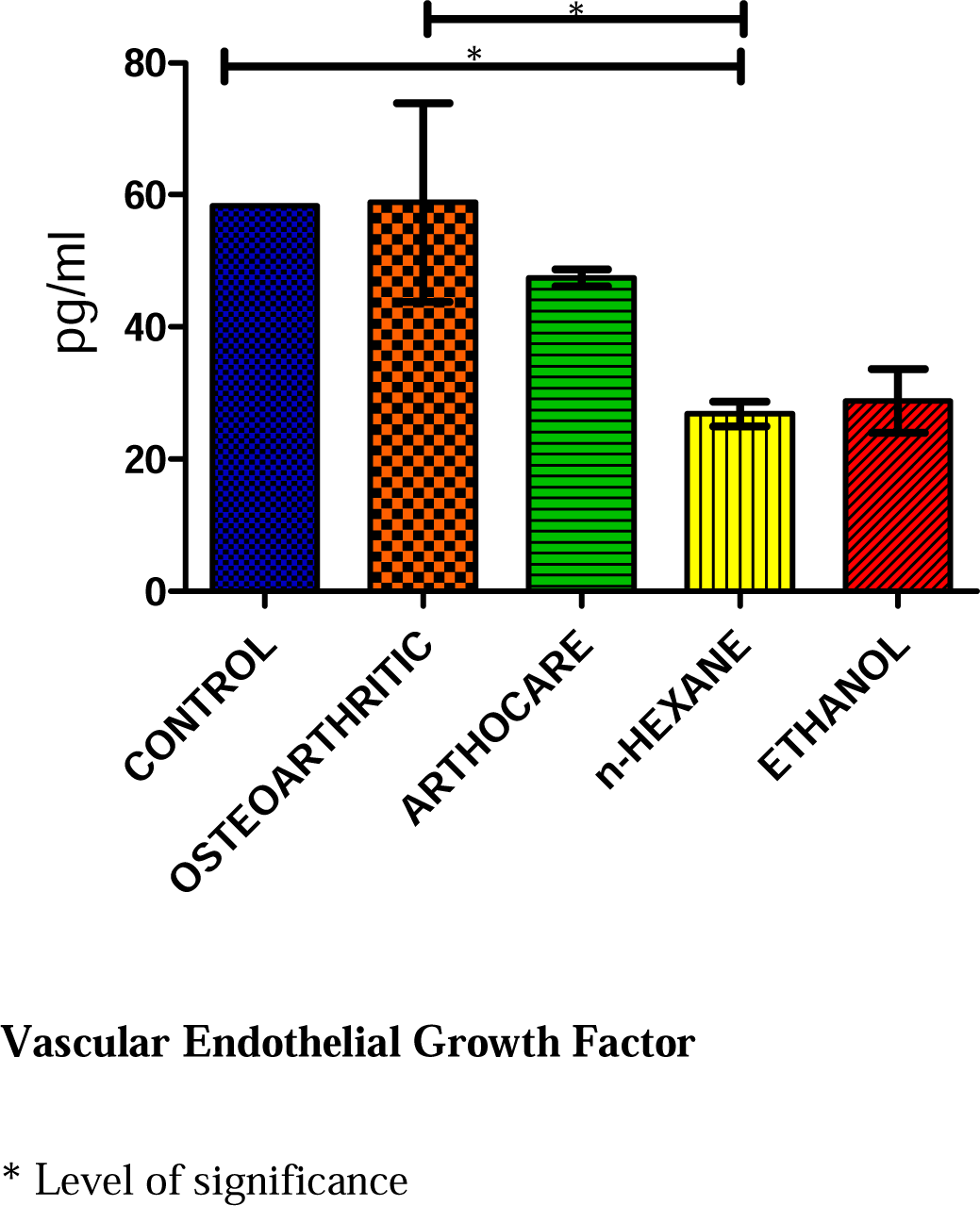

The level of VEGF in cartilage showed significant (p<0.05) decrease when n-Hexane group was compared with control group. There was significant (p<0.05) decrease when n-Hexane group was compared with osteoarthritic group. There was no significant (p<0.05) decrease when n-Hexane group was compared with arthocare group and ethanol group.

The level of VEGF in cartilage showed no significant (p<0.05) decrease when ethanol group was compared with control group, osteoarthritic, arthocare and n-Hexane group.

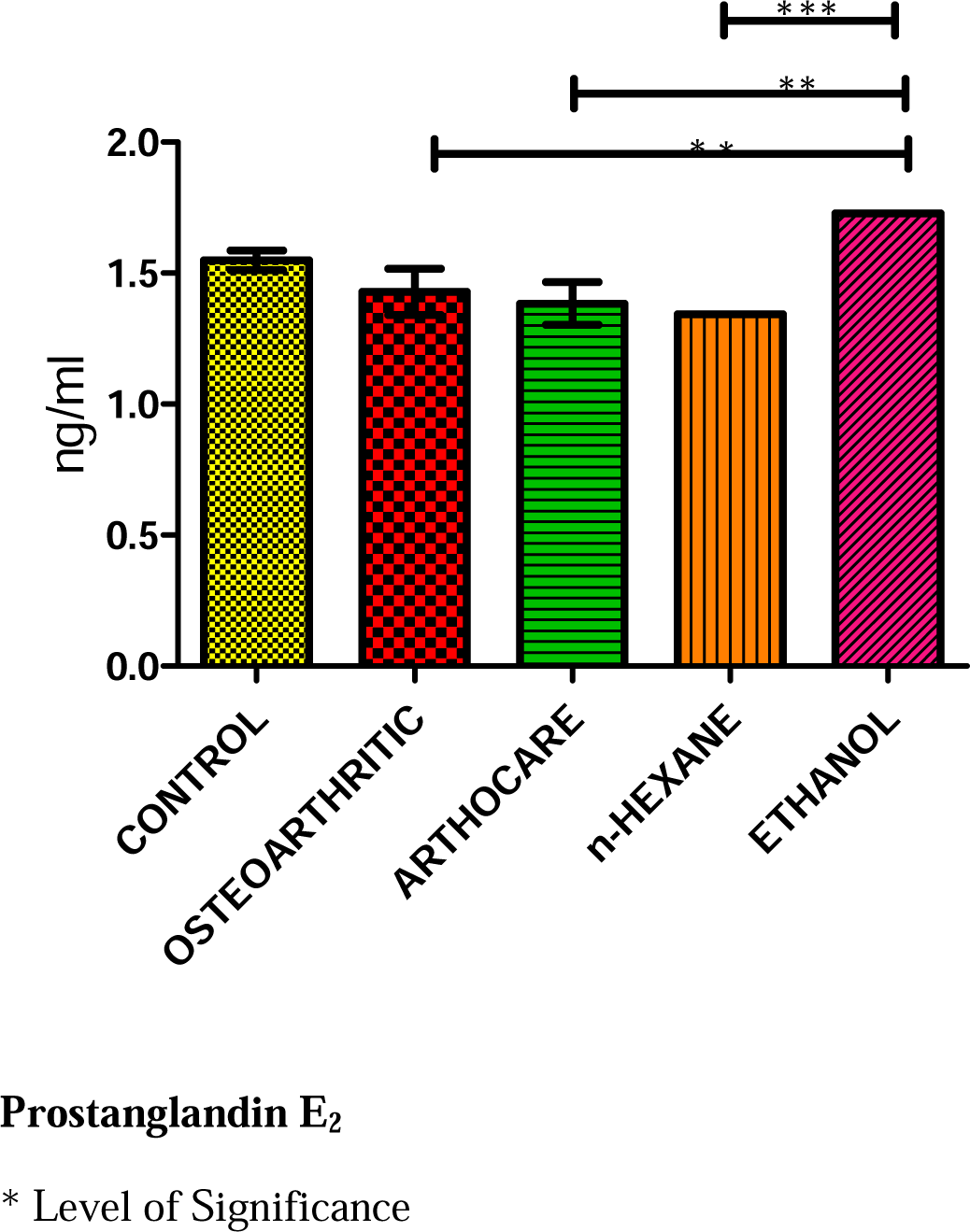

There was no significant (p<0.05) decrease in the level of PGE_2_ in the cartilage when n-Hexane group was compared with control, osteoarthritic, arthocare. There was significant (p<0.05) decrease in the level of PGE_2_ in the cartilage when n-Hexane group was compared ethanol group.

There was significant (p<0.05) increase in the level of PGE_2_ in the cartilage ethanol group was compared with osteoarthritic, arthocare and n-Hexane groups.

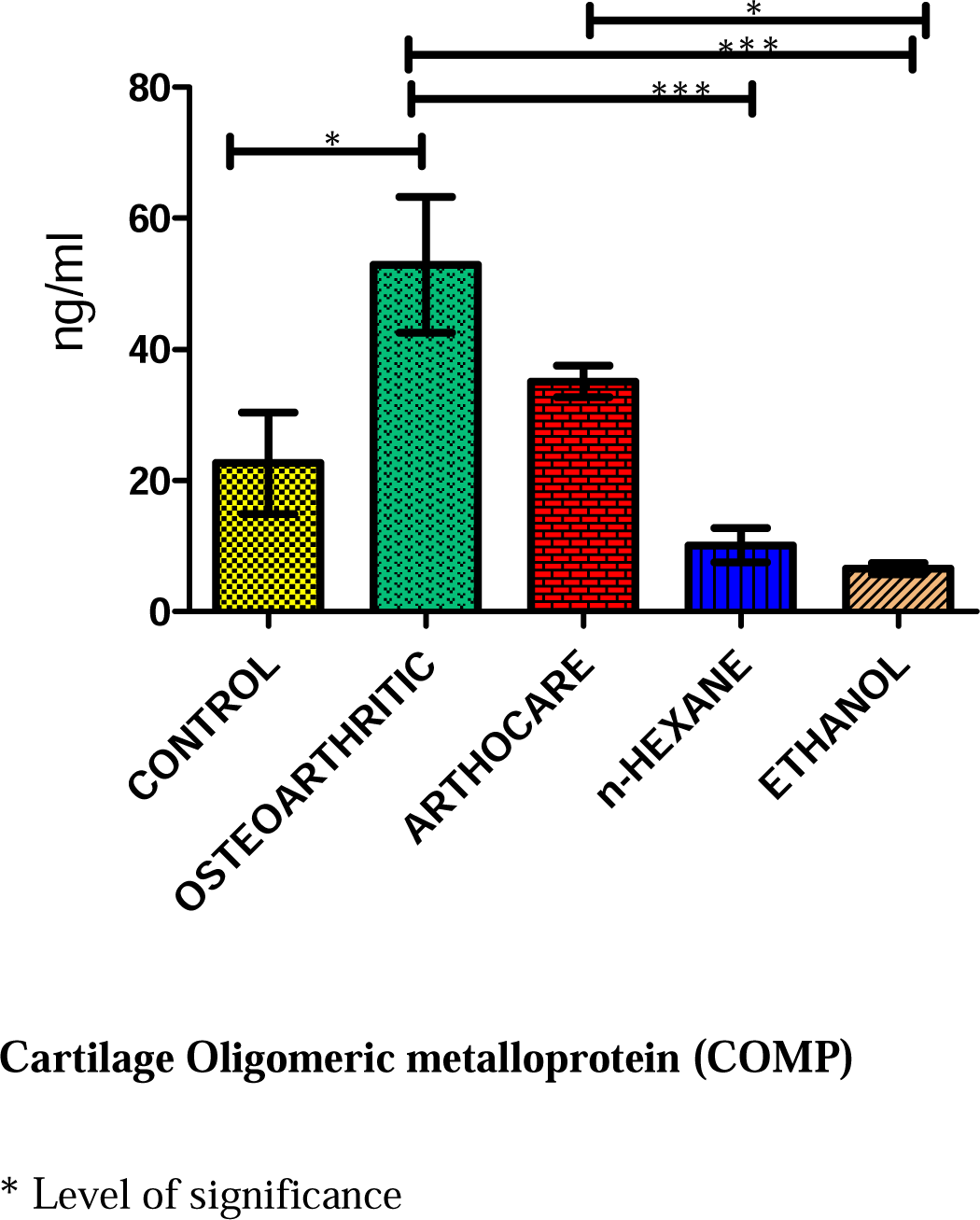

The level of COMP in the cartilage showed no significant (p<0.05) decrease when n-Hexane group was compared with control group. There was significant (p<0.05) decrease when n-Hexane group was compared with osteoarthritic group. There was no significant (p<0.05) decrease when n-Hexane group was compared with both arthocare and ethanol groups.

The level of COMP in the cartilage showed no significant (p<0.05) decrease when ethanol group was compared with control group. There was significant (p<0.05) decrease when ethanol group was compared with both osteoarthritic and arthocare group. There was no significant (p<0.05) decrease when ethanol group was compared with n-Hexane groups.

### KNEE OEDEMA

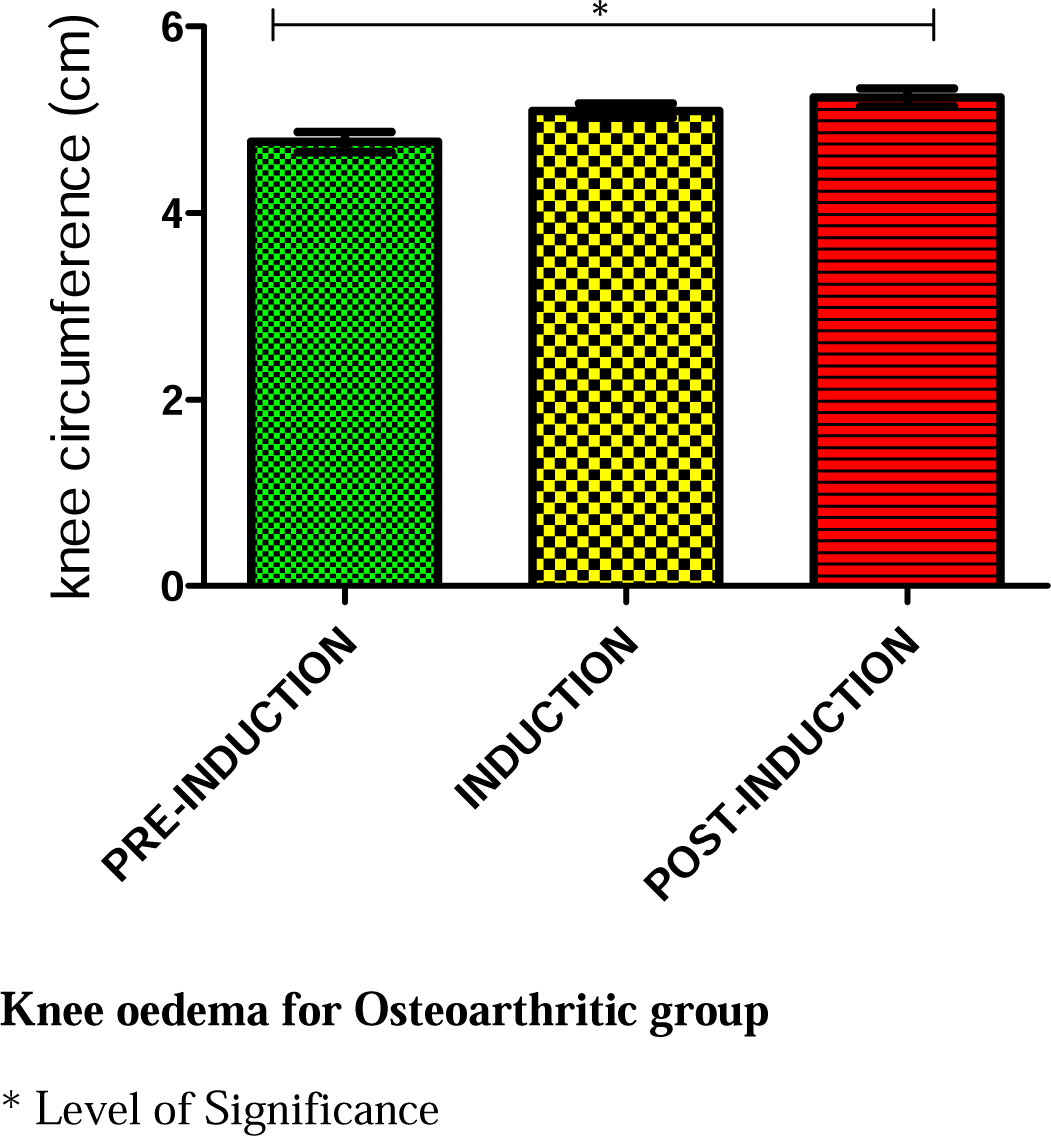

Knee oedema for osteoarthritic group shows significant (p<0.05) difference between pre-induction and post induction.

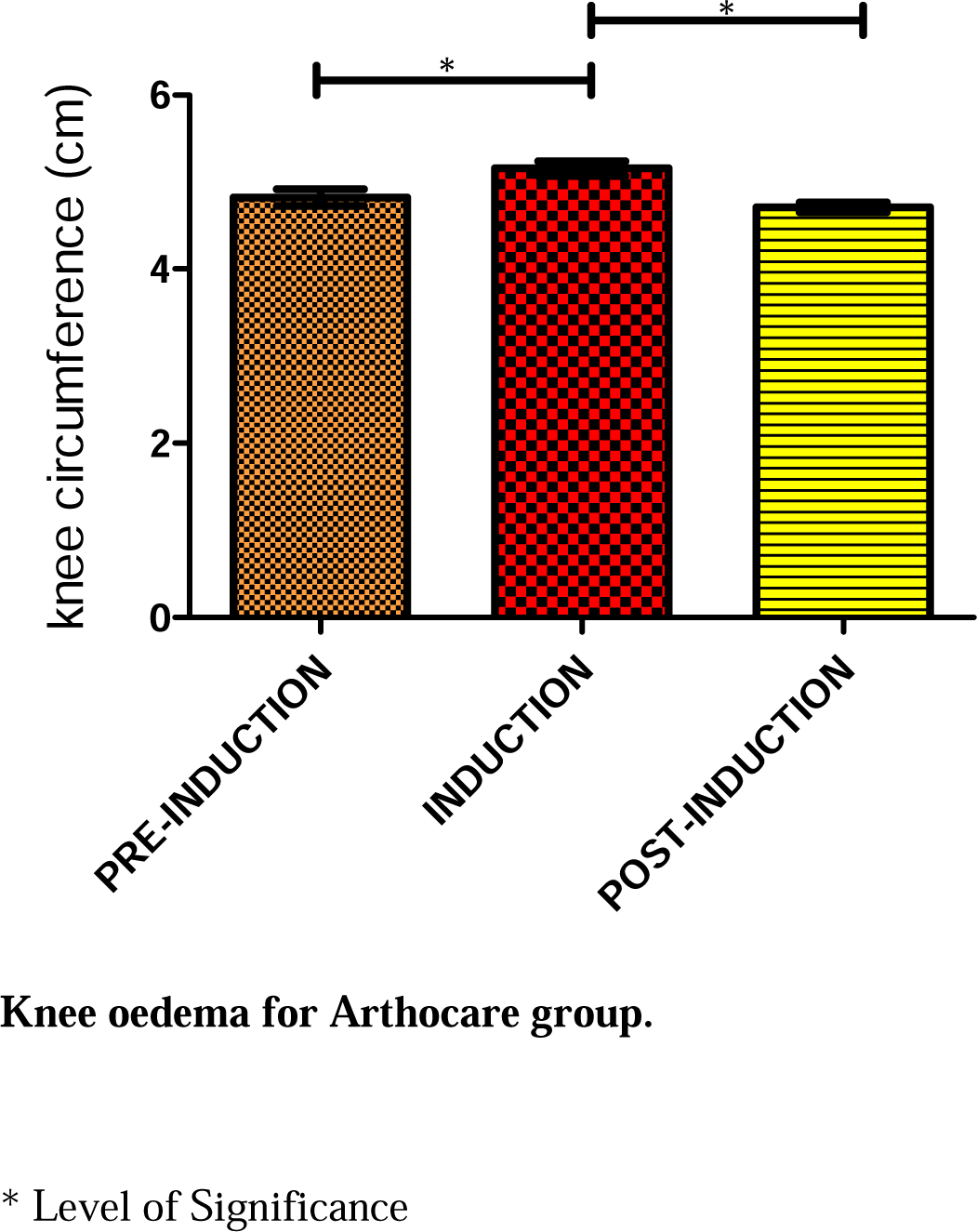

Knee oedema for Arthocare group shows significance (p<0.05) between pre-induction and induction, also between induction and post induction.

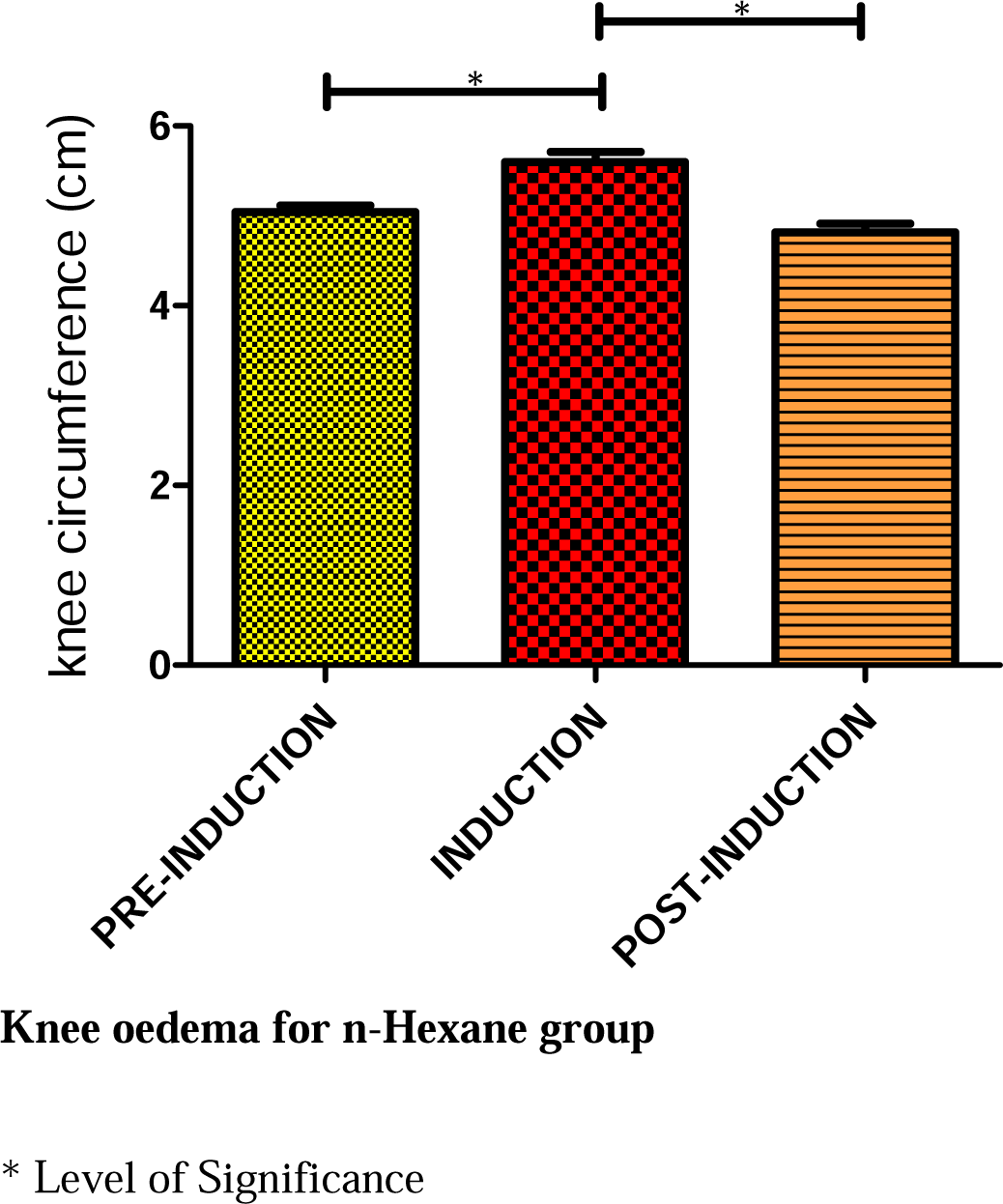

Knee oedema for n-Hexane group shows significance (p<0.05) between pre-induction and induction, also between induction and post induction.

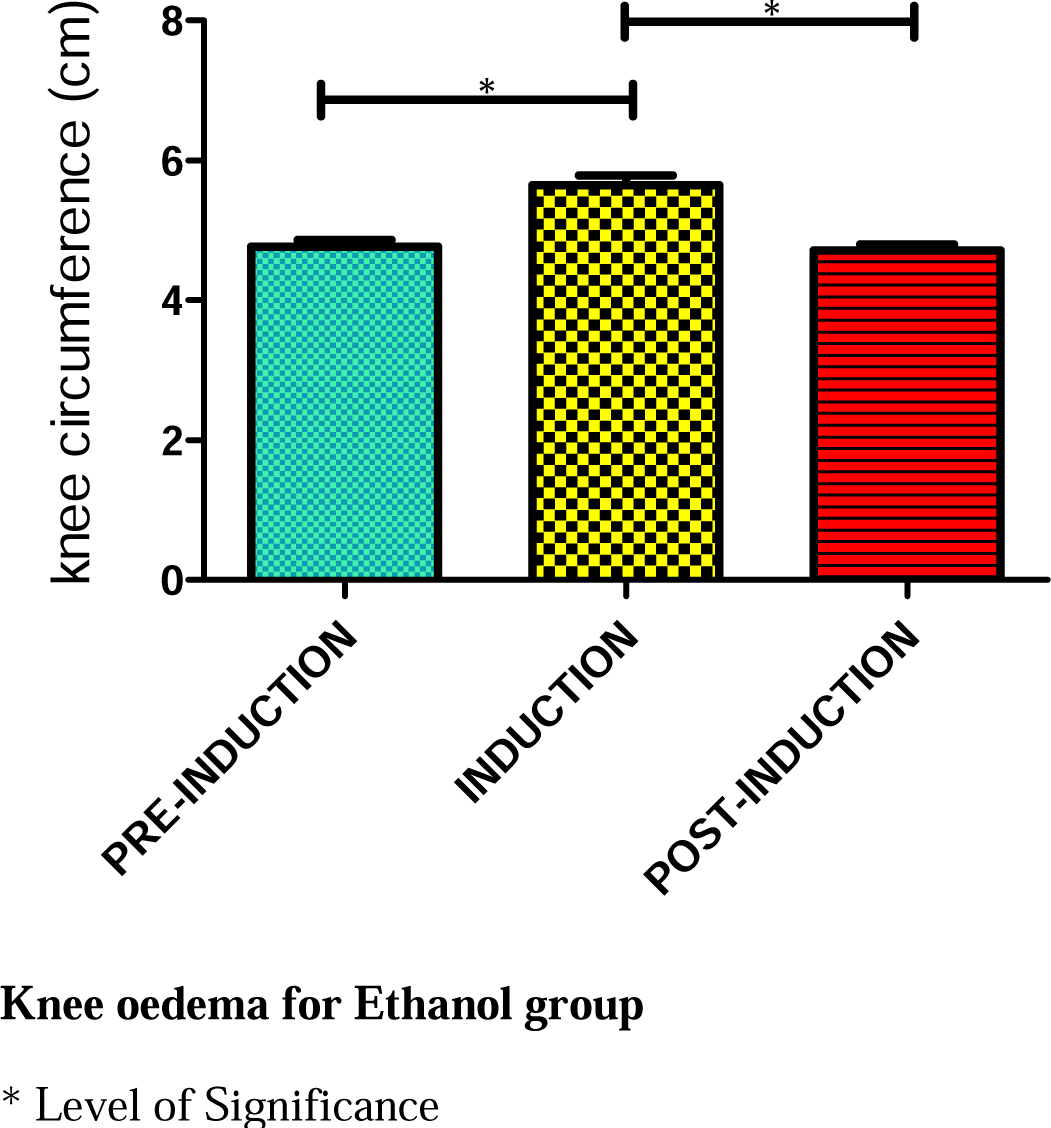

Knee oedema for Ethanol group shows significance (p<0.05) between pre-induction and induction, also between induction and post induction.

### ASSESSMENT OF TACTILE ALLODYNA

#### Mechanical sensitivity

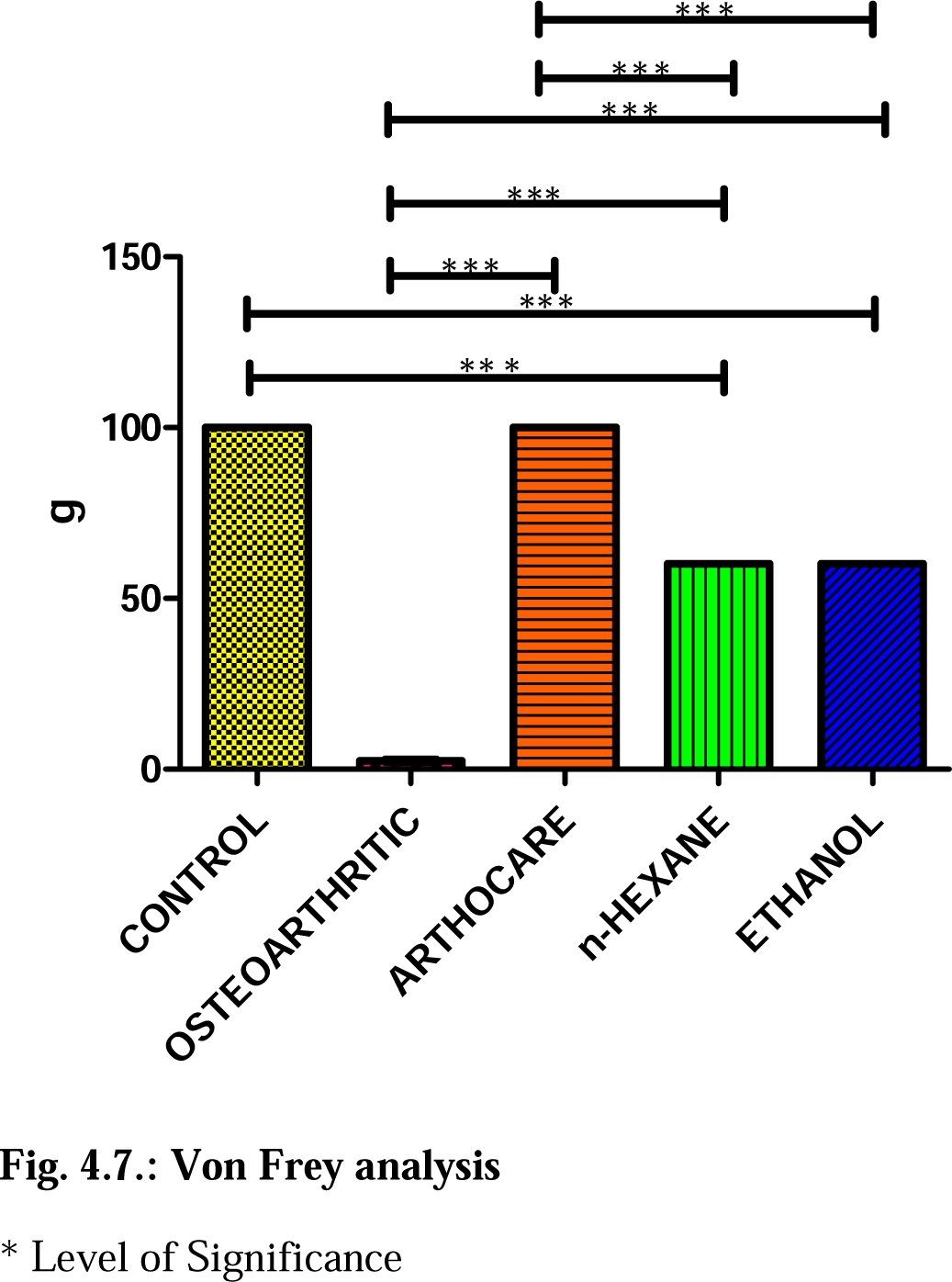

Mechanical sensitivity of n-Hexane group shows significant (p<0.05) increase when compared with Control group, Osteoarthritic group and Arthocare group. But no significance (p<0.05) increase was observed when compared with Ethanol group. Also, the Mechanical sensitivity of ethanol group shows significant (p<0.05) increase when compared with control group, Osteoarthritic group and Arthocare group. But no significance (p<0.05) increase was observed when compared with n-Hexane group.

## DISCUSSION AND CONCLUSION

### DISCUSSION

Over the years, there have been fewer biological experiments performed with the use of *Milicia excelsa* stone. This research was aimed at investigating the biological activity of *Milicia excelsa* on pain perception in female osteoarthritic Wistar rats.

Biomarkers of cell degeneration or inflammation act as prognostic measures of disease progression or treatment outcome in osteoarthritis. These molecules are products of metabolism released into the serum, urine, synovial fluid and their level correlates with osteoarthritic changes in joints. The biomarkers are Tumour Necrotic factor Alpha (TNF-α), and Prostanglandin E_2_ (PGE_2_). Therefore, the development of cartilage pathology in Osteoarthritis involves excessive damage to the collagen fibrillar network, which appears to be mediated primarily by the chondrocyte-generated cytokines IL-1β and TNF-α. However, several biomarker assays have been developed which can be used to measure the synthesis and degradation of collagen fragments, and therefore provide information regarding cartilage turnover.

TNF-α as inflammatory marker plays a major role in pathophysiologic processes of Osteoarthritis. The TNF-α secretion is correlated synergistically with the level of IL-1β. A major risk factor in injuries is mechanical stress which results in the subsequent release of cartilage matrix degradation products during starting and progression of Osteoarthritis. This situation triggers the same signaling pathways as those induced by pro-inflammatory biomarkers including TNF-α and IL-1β. It means that the mentioned cytokines trigger inflammation and catabolic processes in joint tissues such as cartilage and synovial fluids, and so activate inner cell pathways. For example, TNF-α suppresses the syntheses of type II collagen and proteoglycan. Hence, MMP-1, MMP-3, MMP-13, a kind of disintegrin, and metalloproteinase with thrombospondin motifs-4 (ADAMTS-4) are synthesized in induced chondrocytes. It is shown that there is a positive correlation between the expression of TNF receptor and pain, joint stiffness, and disease severity in serum of OA patients. As a result, it is proposed that TNF-α is a good biomarker for OA. COMP also, has shown promise as a diagnostic and prognostic indicator and as a marker of the disease severity as well as the effect of treatment. It is a good prospect for detecting early stage of Osteoarthritis. Enzyme-linked immunosorbent assays (ELISAs) for the detection of this protein and its fragments in synovial fluid and serum have been developed and tested in patients with knee and hip Osteoarthritis and other forms of inflammatory arthritis. Persistently high serum levels COMP have been detected in patients with traumatic knee injury and posttraumatic Osteoarthritis. Large-scale population studies (the Johnston County Osteoarthritis Project) have confirmed that serum COMP protein reflects presence and severity of Osteoarthritis. Substantial amounts of COMP are produced by several mesenchyme-derived cells including synoviocytes and dermal fibroblasts. These findings raise important concerns regarding the utility of measurements of COMP levels in serum or in synovial fluid as markers of articular cartilage degradation because of the likelihood that a substantial proportion of COMP or COMP fragments present in serum or synovial fluid may be produced by cells other than articular chondrocytes.

In addition to all Osteoarthritis biomarkers described above, there are epigenetic biomarkers which have also emerged as a key mechanism in the development of Osteoarthritis. Moreover, Vascular Endothelial Growth Factor **(VEGF)** can be used for determining severity of OA. The expressions of TNF-∞ during osteoarthritis enhance the secretion of Vascular Endothelial Growth Factor (VEGF). The mechanism for this is TNF-∞ enhances induces osteoponin expression (kaomongkolgit *et al.,* 2010). Osteoponin enhances the expression of VEGF through the phosphorylation of AKT and extracellular signal regulated kinase (Dai *et al.,* 2009).

For the purpose of this study, the biomarkers considered were Tissue Necrotic Factor, Prostanglandin E_2_ Vascular endothelial growth factor (VEGF), and Cartillage oligomeric metalloprotein (COMP). There was a significant (p<0.05) decrease in the level of serum Tissue Necrotic Factor (TNF-α) in both n-Hexane and ethanol extract group of *milicia excelsa* stone. This is consistent with the observation of Tsukamoto where he recorded that calcium carbonate decreased secretion of Tissue necrotic factor alpha (Tsukamoto *et al.,* 1996), since *milicia excelsa* stone is rich in calcium carbonate. The result of was also consistent with the effect of the established drug. The mechanism by which this is accomplished is poorly understood.

There was a significant (p<0.05) decrease in the level of Prostanglandin E_2_ in both n-Hexane and Ethanol extract group of *milicia excelsa* stone in the serum. The mechanism for this is proposed to be the reduction in the secretion of Tissue necrotic factor alpha (TNF-α). This is because (TNF-α) as inflammatory biomarker stimulates the secretion of interleukin 18 (il-18). Interleukin 18 stimulates the secretion of Prostanglandin E_2_ (Gogebakan *et al.,* 2016). Therefore decrease in the secretion of TNF-а will reduce Prostanglandin E_2_. But this was not the same for the level of PGE_2_ in the cartilage. There was no significant (p<0.05) decrease in the level of PGE_2_ when n-Hexane group was compared with control, osteoarthritic and arthocare group. The level of PGE_2_ increased in the ethanol group. The mechanism for this increase is poorly understood.

There was a significant (p<0.05) decrease in the level of VEGF both in the serum and cartilage. The mechanism for this is proposed to be due to decrease in TNF-∞ secretion. Since TNF-∞ induces osteoponin and osteoponin enhances the expression of VEGF through phosphorylation (kaomongkolg *et al.,* 2010), a decrease in TNF-∞ will decrease osteoponin secretion which will inturn decrease VEGF secretion.

There was a significant (p<0.05) decrease in the level of COMP both in the serum and cartilage. The mechanism for the action of *Milisia excelsa* stone on COMP is not known.

The results of the biomarkers were also consistent with results of the knee oedema. For n-Hexane and Ethanol group, there was a significant (p<0.05) decrease in the degree of oedema. This decrease is believed to be due to the reduction in cytokine secretion in the joint. In the Osteoarthritic group, there was no significant difference between induction and post induction.

The result of mechanical sensitivity was consistent with those of both biomarkers and knee oedema. For n-Hexane and Ethanol group, there was a significant (p<0.05) increase in their level of mechanical sensitivity when compared with the Control group and Osteoarthritic group. This is also due to reduction in both TNF-α and Prostanglandin E_2._There was no significant difference between n-Hexane and Ethanol group.

Radiographic evaluation showed successful induction of osteoarthritis in the right limb of the female Wistar rats. Also, it showed the effect of n-Hexane and ethanol extract on osteoarthritis.

Although, few researches have been conducted in line with the use of *milicia excelsa* in pain perception, some of the results obtained in this work are consistent with the work of Tsukamoto on the effect of calcium carbonate on pain which is the major component of *milicia excelsa* on TNF-α.

### CONCLUSION

*Milicia excelsa* helps to reduce pain by the effect of calcium carbonate on TNF-α and Prostanglandin E_2_. Both n-Hexane extract and the ethanol extract had the same effect as the established drug, arthocare, which shows a promising area for further research.

## Notes

### Competing Interest Statement

The authors have declared no competing interest.

